# Aβ, tau, α-synuclein, huntingtin, TDP-43, PrP and AA are members of the innate immune system: a unifying hypothesis on the etiology of AD, PD, HD, ALS, CJD and RSA as innate immunity disorders

**DOI:** 10.1101/000604

**Authors:** Claudiu I. Bandea

## Abstract

Despite decades of research, thousands of studies and numerous advances, the etiologies of Alzheimer’s Disease (AD), Parkinson’s Disease (PD), Huntington’s Disease (HD), Amyotrophic Lateral Sclerosis (ALS), Frontotemporal Lobar Degeneration (FTLD-U), Creutzfeldt-Jakob Disease (CJD), Reactive Systemic Amyloidosis (RSA) and many other neurodegenerative and systemic amyloid diseases have not been defined, nor have the pathogenic mechanisms leading to cellular death and disease. Moreover, the biological functions of APP/amyloid-β (Aβ), tau, α-synuclein, huntingtin, TAR DNA-binding protein 43 (TDP-43), prion protein (PrP), amyloid A (AA) and some of the other primary proteins implicated in amyloid diseases are not known. And, there are no successful preventive or therapeutic approaches. Based on a comprehensive analysis and new interpretation of the existing data in context of an evolutionary framework, it is proposed that: (i) Aβ, tau, α-synuclein, huntingtin, TDP-43, PrP and AA are members of the innate immune system, (ii) the isomeric conformational changes of these proteins and their assembly into various oligomers, plaques, and tangles are not protein misfolding events as defined for decades, nor are they prion-replication activities, but part of their normal, evolutionarily selected innate immune repertoire, and (iii) the immune reactions and activities associated with the function of these proteins in innate immunity lead to AD, PD, HD, ALS, CJD, RSA and other related diseases, which are innate immunity disorders.

---

Despite decades of research, thousands of studies and numerous advances, the etiologies of Alzheimer’s Disease (AD), Parkinson’s Disease (PD), Huntington’s Disease (HD), Amyotrophic Lateral Sclerosis (ALS), Frontotemporal Lobar Degeneration (FTLD-U) and Creutzfeldt-Jakob Disease (CJD), and many other neurodegenerative and systemic amyloid diseases have not been defined, nor have the pathogenic mechanisms leading to cellular death and disease. Moreover, the biological functions of APP/amyloid-β (Aβ), tau, α-synuclein, huntingtin, TAR DNA-binding protein 43 (TDP-43), prion protein (PrP) and some other primary proteins implicated in amyloid diseases are not known. And, there are no successful preventive or therapeutic approaches.

Based on a comprehensive analysis and new interpretation of the existing data, it was recently proposed that Aβ, tau, α-synuclein, huntingtin, TDP-43, PrP, and other primary proteins implicated in neurodegenerative disorders are members of the innate immune system, and that their “malfunction” leads to a wide range of autoimmune disorders, including AD, PD, HD, ALS, FTLD-U, and CJD (1,2). It is likely, also, that other amyloid-forming proteins, such as amyloid A (AA) which is implicated in Reactive Systemic Amyloidosis (RSA (3,4), belong to this group of innate immunity proteins. Here, I present evidence and arguments supporting this unifying hypothesis, which, if correct, would radically change the understanding of these devastating diseases and help with the development of preventive, diagnostic, and therapeutic approaches.

## Aβ, tau, α-synuclein, huntingtin, TDP-43, PrP and AA are members of the innate immune system

According to this unifying hypothesis, Aβ, tau, α-synuclein, huntingtin, TDP-43, PrP, and AA perform their innate immunity function by participating in two major, overlapping mechanisms or pathways: (i) blocking the life cycle of various microbial and viral pathogens directly, for example, by damaging the microbial cellular membrane or the host cell membranes required for viral replication, or (ii) indirectly, inducing the death of host cells by various non-inflammatory or inflammatory mechanisms, including apoptosis, which limit or block the spread of infection. Additionally, these innate immunity proteins exercise their protective functions in other types of injuries that mimic those produced by infectious pathogens, including physical, biochemical, immunological, and age-related injuries or dysfunctions.

To perform their protective roles, these putative innate immunity proteins can assume multiple functional isomeric conformations, and can assemble into diverse functional oligomeric structures, or *innate immunity complexes* (IICs). In their native, ‘unengaged,’ ligand-like conformation, these proteins can exist as monomers or small oligomers that are relatively unstructured and contain primarily α-helix protein folds. Upon contact with microbial or viral components, or after detecting ‘danger signals’ associated with diverse microbial and viral infections or other types of injuries, these relatively unstructured proteins assemble into a population of diverse IICs by assuming new isomeric conformations that are rich in β-sheet folds, a defining characteristic of amyloids. These IICs range in size and shape, from relatively small globular oligomers to larger amyloidal fibrils, tangles, plaques, and amyloid deposits.

To be able to interact with various microbial and viral pathogens or to signal infection and amplify the signals associated with infections, these innate immune proteins can assemble into *pathogen-like IICs* (PICs) that mimic or simulate microbial or viral molecular patterns or activities. For example, some of these proteins can form channels or pores (reviewed in 5,6; see below) that damage the membrane of microbial pathogens or the host intracellular membranes associated with viral replication. However, by mimicking or simulating the components or activities of various pathogens, some of these putative innate immunity proteins can also assemble into *toxic PICs* (tPICs) that induce or signal the death of the infected cells and tissues by various mechanisms. Although destructive at the cellular or tissue level, by blocking the spread of infections, these *immune pathogenic reactions* (IPRs) are protective at the individual and population level.

The innate immunity pathways described above are consistent with the vast number of data and observations regarding the expression patterns and the properties of this group of putative innate immune proteins (see discussion in 2). The ability of many amyloid proteins, including Aβ, α-synuclein, huntingtin, PrP, and AA, to assemble into membrane-damaging channels or pores has been documented in hundreds of studies (reviewed in 5-7). Also, there are direct experimental data and observations supporting the antimicrobial and antiviral activities of this group of putative innate immunity proteins (8-17) and the interaction of these proteins with diverse infectious agents or with various arms of the immune system (e.g., see 18-33; note: though not recognized as such, the RNAi system is an important arm of the innate immune system; 5,34.) Although many of these studies were designed to address the pathogenic mechanisms by which these proteins cause disease, not their physiologic function, their results are nonetheless consistent with and support the new unifying hypothesis.

In addition to these innate immunity proteins, many well-known members of the immune system, including the interferon- and TNF-families (36,37; discussed in 1), and newly discovered members such as mitochondrial antiviral signaling protein (MAVS) (38-40), assemble into pathogen-like amyloid aggregates in order to perform their innate immunity function. Moreover, many genuine antimicrobial proteins perform their innate immunity functions by assembling into “toxic” pores (reviewed in 41) or, as is the case with protegrin-1 (42), by assembling into amyloid fibers. Interestingly, the RNA-dependent RNA polymerase of many RNA viruses, including polioviruses and other neurotropic viruses, assembles into amyloid-like oligomers, lattices, and tubular formations rich in β-sheet domains on the surface of various intracellular membranes (e.g., mitochondria), where they direct the synthesis of viral RNA genome and transcripts (43,44). It is easy to envision how, by assembling into membrane-associated IICs, some of these putative innate immunity proteins, such as Aβ, can disrupt the life cycle of these viruses (discussed in 2).

Interestingly, both Aβ and PrP share structural and sequence domains with viruses (45-52), which suggests not only a potential antiviral immune function, but also an endosymbiotic viral evolutionary origin (1,2; see below). In turn, many microbial and viral pathogens employ a wide range of immune silencing mechanisms that are based on products that structurally or functionally mimic the components and activities of the immune system (53,54).

As outlined above, this group of putative innate immunity proteins is also involved in protecting against non-microbial or non-viral pathogenic products resulting from physical, biochemical, and age-related injuries or dysfunctions, particularly those that resemble products and activities associated with microbial or viral pathogens. However, the most remarkable tenet of this hypothesis, which is key to understanding the etiology of these neurodegenerative and systemic amyloid disorders, is that these innate immunity proteins are also critical in conferring “protection” against some of their own IICs, particularly the tPICs which are involved in *immune pathogenic reactions* (IPRs), by sequestering them into protective IICs in the form of large oligomers, fibers, plaques, and amyloid deposits.

## The cycling autoimmune reactions associated with the function of amyloidogenic proteins in innate immunity lead to AD, PD, HD, ALS, FTLD-U, CJD and RSA

The molecular mechanisms employed by Aβ, tau, α-synuclein, huntingtin, TDP-43, PrP, and AA to assemble into diverse IICs are based on their essential properties to: 1) recognize viral- or microbial-like molecular patterns, and 2) assemble into IICs that mimic or simulate these molecular patterns, the PICs. In turn, the PICs become targets for the ‘parental’ native proteins, which assemble into additional PICs in a series of *cyclic autoimmune reactions* (CARs), which amplify the immune response. The CARs are resolved by the seclusion of PICs into larger IICs in the form of plaques, tangles, and amyloid deposits. Additionally, these innate immunity proteins work in concert with other arms of the immune system and with the protein metabolic processes, such as autophagy (e.g., see 55,56), in resolving the CARs and clearing the IICs and the associated microbial, viral and host cellular debris.

Although, similar to other members of the immune system, these putative innate immunity proteins have been strongly selected against extended pathogenic reactions that would lead to autoimmune diseases, they run a fine line between ‘protection’ and ‘pathogenicity.’ The line is particularly thin in this group of proteins because, to protect the organism at the individual level, some of the IICs (i.e., tPICs) cause the death of infected cells and tissues by various IPRs. Moreover, the CARs are based on the property of the native isomeric conformers of these proteins to recognize the PICs and assemble into new PICs, which fundamentally is an ‘autoimmune reaction.’ As previously proposed (1,2), some of these proteins, such as PrP, have the extraordinary property to recognize and fold into many isomeric conformations, which enable them to interact with numerous pathogens and provide a multivalent immune response (interestingly, it was recently shown that PrP interacts with various pathogen-like, β-sheet-rich conformers independent of their amino acid sequence; 57.) This remarkable isomeric flexibility confers upon PrP the potential to assemble into a population of diverse PICs. The native PrP isomers recognize these PICs with differential kinetics, which leads to a selection process and an increase in the efficiency of CARs. Although powerful, this selection-based immune mechanism, which can be regarded as an ‘adaptive immune feature,’ opens the door for pathogenic autoimmune reactions and disease. Indeed, due to mutations or other genetic abnormalities (e.g., gene or chromosomal duplications) that change the amino acid sequence or the expression level of these putative innate immunity proteins, or due to many other risk factors such as persistent or recurrent infections or physical, biochemical, and age-related injuries, these proteins assemble into PICs that become preferred targets for the parental isoforms. Although rare events, the assembly of these PICs, suggestively labeled here *disease-driving PICs* (dPICs), leads to endless *disease-driving CARs* (dCARs), and eventually to AD, PD, HD, ALS, FTLD-U, CJD, and other neurodegenerative or systemic amyloid disorders.

The genes coding for APP/Aβ, tau, α-synuclein, huntingtin, TDP-43, and PrP are expressed primarily in tissues and organs that are not under full surveillance by the adaptive immune system, such as the brain and testes, which supports the unifying hypothesis about their function in innate immunity (note: the relatively high expression of some of these proteins in germline tissues supports the hypothesis that they also play a critical role against viral endogenization and against the spread of endogenous viral elements; discussed in 1,2.). Although the assembly of the incipient dPICs might originate at a higher frequency in these particular tissues, they can assemble *de novo* in all tissues where these innate immunity proteins are expressed. Once formed, however, the dPICs become preferred targets of the ‘parental’ or other proteins from this group of putative innate immunity proteins leading to dCARs and to the production of a large population of additional dPICs.

In addition to the putative selection process involved in increasing the efficiency of CARs, the population of diverse PICs, including dPICs, enters a second selection process— that regarding the ability to circulate among neighboring cells and tissues where they become targets of the resident ‘parental’ innate immune proteins, leading to an *expanded autoimmune cycle* (EAC). Unlike most tissues and organs, in which, due to high cellular turnover, the pathology associated with EACs is limited, in tissues that contain post-mitotic cells, such as the central nervous system, the EACs lead to massive cellular dysfunctions, cellular death, and to a wide spectrum of clinical neurodegenerative diseases.

In rare circumstances, the dPICs can be transferred by various routes, such as food consumption or fluid transfer, between individuals of the same or different species, a phenomenon that resembles the transmission of viruses and other infectious pathogens. Although transmission of amyloidosis among individuals and species has been known and studied for a long time (58,59; see also 4), in the TSE field this phenomenon led to the formulation of the prion hypothesis, which asserts the existence of self-replicating, protein-only infectious agents called ‘prions’ (reviewed in 60). The prion hypothesis has been increasingly associated with other neurodegenerative and systemic amyloid disorders (61-67), which have been traditionally classified as protein misfolding disorders. These two highly influential working hypotheses - the ‘protein misfolding concept’ and ‘prion hypothesis’ - have directed much of the thinking and research during the last few decades and, therefore, in order to be viable, this radical new perspective must explain the data, observations, and paradigms associated with these dogmas.

## By coupling physiology and pathology the unifying hypothesis challenges the protein misfolding concept and the prion hypothesis

Whether they were regarded as the cause or result of disease, the amyloids have been central to the effort to understand the etiology of amyloid disorders, such as AD, for more than a century (68,69). After the discovery that the amyloid aggregates contain primarily polypeptides that acquire a β-sheet-rich structure, the amyloids were classified as “misfolded proteins” and the associated diseases as “protein misfolding disorders” (reviewed in 70).

The classification of the amyloidogenic process as a protein misfolding event has uncoupled the process, its products (i.e., amyloids), and the associated diseases from the physiological function of these proteins, and it has turned the balance in favor of the more intuitive and parsimonious perspective that amyloids are the cause, rather than the result of disease. And, the fact that many of these amyloid-forming proteins were attributed hypothetical functions, such as in metal-ion transport or in neurotransmission, which were not immediately suggestive of a need for the amyloidogenesis process, did not help; it is becoming increasingly evident, however, that amyloids can perform many important biological functions (71-74).

Nevertheless, the critical development that has reinforced the protein misfolding dogma and set a strong conceptual barrier between the physiological function of these proteins and the disease mechanisms has been the prion hypothesis. By defining and promoting the ‘prions’ as novel, protein-only infectious pathogens that self-replicate independently of the PrP gene (60), the prion hypothesis has uncoupled the transmissible spongiform encephalopathies (TSEs) phenomena from the function of PrP and its gene. However, there is substantial genetic, structural, functional, and evolutionary evidence suggesting that the prion hypothesis and the associated concepts of ‘self-replicating proteins’ or ‘prion replication’ are flawed.

First, current evidence indicates that the gene coding for PrP is a symbiotic endogenous viral gene (1,2,75). In addition to the fact that PrP has viral properties and assembles into virus-like structures (for illustrative data on PrP assembly into virus-like structures, see 52; and for the presence of virus-like structures in TSE tissues, see 76-78), the gene coding for PrP shares sequence domains with the retroviral reverse transcriptase gene (49) and with the HIV-1 fusion peptide (51). It appears, also, that the PrP gene codes not only for PrP, but also for TAR-like RNA elements (discussed in 2; for the original data, see 79-81) and for an out-of-frame polypeptide (82), which are features characteristic of viral genes. Thus, from an evolutionary perspective, it appears that the TSEs have an endogenous viral etiology and, therefore, the prion hypothesis, which was specifically proposed and promoted (60) as an alternative to the conventional hypothesis that TSEs have a viral etiology, is questionable.

Second, there is persuasive evidence that PrP is a member of the innate immunity system and that TSEs are autoimmune diseases; thus, from a biological perspective, what has been mistakenly regarded as the ‘prion replication’ phenomena, which challenged the central dogma of molecular biology, are in fact a series of cyclic autoimmune reactions. Moreover, because the prion hypothesis was formulated and promoted based on tenets that circumvented some of the fundamental principles of protein biochemistry and those governing the relationship between genetic information and phenotype (discussed in 1,2), other novelties associated with the prion hypothesis such as the ‘prion hereditary information’ and ‘prion strains’ might be flawed.

It is well known that the vast majority of TSE cases start spontaneously, in the absence of previous ‘prions,’ which means that the ‘prions’ can arise without the so-called ‘prion hereditary information.’ Also, it has been known for a long time (however, see 83) that, just like PrP, thousands of other proteins undergo and achieve their various transitional and functional isomeric conformations, and that they assemble into various protein complexes by using not only genetic information (i.e., the nucleic acid-based information coding for the order of amino acids in the polypeptide chains), but also biological information encoded in many other molecules, which are used as chaperones, templates, partners, substrates, etc. Indeed, similar to the assembling process of PrP into various IICs, including dPICs, many other proteins, some of them members of the immune system, undergo various isomeric transitions and interact with other protein molecules in order to assemble into functional complexes, including amyloid fibrils. The MAVS is an illustrative example of an innate immune protein that, similar to PrP, assembles into fibers in order to achieve its antiviral isomeric conformation (38) and, apparently, the macrophage migration inhibitory factor also assembles in amyloid fibrils in order to perform its immunological function (84). Moreover, the assembly of thousands of proteins into dimers, oligomers, or larger complexes is a ‘protein-only’ affair. Ironically, there’s strong evidence that this is not the case with the PrP, which apparently requires small nucleic acids (85; reviewed in 1,2) for efficient assembly into the so-called ‘prions’; obviously, the participation of nucleic acids in the assembly of the ‘prions’ clashes with the ‘protein-only’ paradigm and with the prion hypothesis.

During CARs, the PrP and other members of this group of putative innate immunity proteins can assemble into PICs that resemble, or are identical to the targeted PICs, giving the false impression of ‘prion replication,’ ‘prion hereditary information,’ and ‘prion strains.’ A related, and one of the most intriguing phenomena in the TSE field, which has received relatively little attention (however, see 86,87) because it is difficult to explain in the context of the prion hypothesis (discussed in 88), is the decoupling of prions’ infectivity/replication from the associated toxicity/pathogenicity. According to the current data (86,87), the ‘infectious’ and the ‘toxic’ forms of PrP are not the same entities, which brings into question the definition of ‘prions’ (88) and has significant implications for understanding the pathogenesis of TSEs and the development of effective therapy (86,87,89). These intriguing phenomena are explained by the CAR-associated selection processes discussed in the previous section, which lead to the production of two major overlapping populations of dPICs: those that circulate with high efficiency and those that are primarily associated with pathogenicity.

Although an interesting biological phenomenon and, certainly, of great public health concern, the transfer of TSE-associated dPICs from one individual or species to another is a relatively rare event. In the vast majority of TSE cases, and in the other neurodegenerative disorders, the incipient dPICs originate *de novo*. As outlined above, the assembly of Aβ, tau, α-synuclein, huntingtin, TDP-43, and PrP into dPICs is a rare, inadvertent event that can be enhanced by various genetic, physiologic, and environmental factors, including chemical, biochemical, and physical injuries, as well as by various infectious agents and diseases. Some of these factors might also play a significant role in the circulation of dPICs among cells and tissues. Noticeably, many of these factors have been defined as risk factors for this group of neurodegenerative diseases, which has led to numerous, divergent hypotheses about the causes and the etiologies of these diseases.

## The unifying hypothesis integrates many current views and paradigms

The strength of the present hypothesis is not only that it is consistent with the large amounts of data, and it has considerable explanatory power, but that it unifies many of the current, often conflicting views regarding the etiologies of these devastating diseases. In the AD field, for example, this hypothesis solves one of the most disputed issues, that regarding the ‘pathogenic’ vs. ‘protective roles of Aβ and its various IICs, which has been highlighted in dozens if not hundreds of publications (e.g., see 56,68,69,90-93); same is true for other proteins in this group (e.g., see 24,94,95). According to this new, unifying hypothesis, which integrates both perspectives, the Aβ and its IICs can have both ‘protective’ or ‘pathogenic’ attributes, whether their activity is defined at the cellular/tissue level or at the organism level.

Among the major risk factors associated with neurodegenerative disorders are: (i) specific mutations in the genes coding for the primary proteins implicated in the diseases, or in the genes coding for other proteins that are implicated in their metabolism or that interact with these primary proteins, and (ii) the aging process. The disease cases associated with mutations in the germline are referred to as hereditary, or familial, and the others, which are the vast majority, are classified as spontaneous. However, due to numerous mutations occurring in somatic cells, it is likely that many of the sporadic cases start with dPICs that originate in cells carrying somatic mutations in these genes (for a documented example in a CJD case, see 96). Either way, by changing the expression level of these proteins, or their amino acid sequence, these mutations, as well as many of the other risk factors implicated in these disorders, increase the odds for the inadvertent formation of the incipient dPICs. Also, within the framework of this unifying model, it is expected that the formation of dPICs and the rate of dCARs are influenced by infectious agents such as herpes simplex virus, by biochemical events such as oxidative stress, or by physical traumas that mimic infection-associated pathogenic events, all of which have been proposed as hypothetical etiologies for these disorders. And, because the aging process is a confounding aspect of these risk factors, and because the EAC is a relatively lengthy process in itself, age becomes an integral facet of many of these neurodegenerative disorders (for a recent study evaluating the length of the pathogenic processes in AD, see 97).

Another integrative paradigm associated with this model is that concerning the wide range of IICs in regard to their structure and size (i.e., from various oligomers, e.g., see 98, to large plaques and tangles), composition (i.e., one or more protein species), and in regard to their ‘pathogenic’ or ‘protective’ effects, which together add a whole new level of integrative and explanatory power to this model. Also, according to this hypothesis, some of the members of this group of putative innate immunity proteins interact with each other during both their normal protective immune functions and during neurodegeneration. For example, the dPICs and some of the other IICs can become targets, or they can signal an innate immune response by other members of this group of innate immunity proteins (57,99). This unifying hypothesis explains the numerous findings suggesting various direct and indirect interactions among the members of this group of proteins. Moreover, the IICs can become targets not only for this group of innate immunity proteins, but also for other components of the immune system. Also, it is important to realize that even the protective IICs, such as the plaques, could interfere with the normal biochemical, physiological, and immune activities, thereby potentially enhancing the pathology associated with these diseases.

## Validation

The hypothesis outlined here is supported by direct evidence, it is consistent with the vast amounts of experimental data and observations, and it makes biological and evolutionary sense. Moreover, this hypothesis explains many enigmatic features associated with neurodegenerative diseases and integrates many of the current ideas and hypotheses on the etiology of these disorders. However, this hypothesis needs full experimental evaluation.

The function of Aβ, α-synuclein, tau, huntingtin, TDP-43, PrP, and AA in innate immunity can be addressed by studying the susceptibility of animal models or of individuals (in natural populations) with differential expression of these putative innate immunity proteins to various pathogens, particularly to neurotropic infectious agents. These studies are facilitated by the existence of numerous transgenic animal models, and by well-defined molecular markers and powerful assays. These studies can be paralleled by cell culture and *in vitro* molecular approaches that investigate the interactions of these innate immunity proteins with the life cycle of various pathogens and their components and activities.

The studies addressing the ‘auto-immune’ model are more involved, as they need to demonstrate the assembly of various IICs, particularly dPICs, and the occurrence of dCARs and EACs. They need to address the specific molecular mechanisms leading to cellular dysfunction, cellular death, and neurodegeneration, which overlap with those responsible for their protective innate immune response.

## Preventive, Diagnostic, and Therapeutic Implications

The full range of preventive, diagnostic, and therapeutic implications of this hypothesis, if validated, remains to be explored. However, this hypothesis supports many of the current ideas and approaches on prevention, diagnosis, and therapy. For example, it would make sense to reduce risk factors, such as infections, oxidative stress, or physical trauma, in order to prevent the formation dPICs and reduce the rate of CARs and EACs. From a diagnostic and therapeutic perspective, it would make sense to define and eliminate the dPICs, or block their participation in dCARs, or possibly to stimulate the formation of protective IICs, such as plaques and amyloid deposits.

## Perspective

The unifying hypothesis presented here is a radical departure from the thinking and the working hypotheses that have directed most of the research in the field in the last few decades. Nevertheless, this unifying theory is based on the same experimental data and observations as the current working hypotheses. The difference is in the interpretation of the data. Unlike the current working hypotheses, which are reductionist in nature, the new theory integrates the vast amounts of experimental data and observations in a comprehensive conceptual framework that make evolutionary and biological sense and has high explanatory power.

Many effective therapies and drugs have been discovered fortuitously, or with little science backup. Although this door of discovery remains open, the chances for it to happen after decades of intense research by both academia and pharmaceutical industry are remote. Therefore, it is likely that the development of preventive and therapeutic approaches requires scientifically sound working hypotheses and, as suggested here and elsewhere (e.g. 100), the current working hypotheses in the field might be flawed.

In light of the extraordinary medical, social and economic burden associated with neurodegenerative diseases, including tens of millions of patients and affected families and huge economic losses estimated at over two hundred billions dollars per year in the Unites States alone, it would make sense, from a scientific and professional perspective, that all new hypotheses in the field are timely, openly and fully evaluated.

